# SNPTB: nucleotide variant identification and annotation in *Mycobacterium tuberculosis* genomes

**DOI:** 10.1101/227066

**Authors:** Aditi Gupta

## Abstract

**Summary:** Whole genome sequencing (WGS) has become a mainstay in biomedical research. The continually decreasing cost of sequencing has resulted in a data deluge that underlines the need for easy-to-use bioinformatics pipelines that can mine meaningful information from WGS data. SNPTB is one such pipeline that analyzes WGS data originating from *in vitro* or clinical samples of *Mycobacterium tuberculosis* and outputs high-confidence single nucleotide polymorphisms in the bacterial genome. The name of the mutated gene and the functional consequence of the mutation on the gene product is also determined. SNPTB utilizes open source software for WGS data analyses and is written primarily for biologists with minimal computational skills.

**Availability and implementation:** SNPTB is a python package and is available from https://github.com/aditi9783/SNPTB

**Supplementary information:** Tutorial for SNPTB is available at https://github.com/aditi9783/SNPTB/blob/master/docs/SNPTB_tutorial.md

## Introduction

Whole-genome sequencing (WGS) has become an
 integral part of basic and translational research on *Mycobacterium tuberculosis,* spanning topics ranging from pathogen evolution, epidemiology, and the emergence of drug resistance to TB latency and vaccine development^1-6^. Publicly accessible databases contain WGS data from thousands of clinical isolates of *M. tuberculosis*^2, 7, 8^. Phenotypic data such as drug susceptibility testing accompanies the WGS for several of these isolates, highlighting the usefulness of this data resource to the scientific community. In particular, identifying single nucleotide polymorphisms (SNPs) in WGS allows determining the functional significance of these mutations, such as their role in the emergence of drug resistance and adaptation to a new environment. In the absence of an open-source and stand-alone computational tool for easy analyses of WGS data, the data accumulation will continue to outpace extraction of meaningful information from sequencing data.

SNPTB is a bioinformatics pipeline that combines widely used open-source software for WGS data quality control, read mapping to the *M. tuberculosis* reference genome, and SNP calling into a single python package. The input to the pipeline is the sequencing data in the fastq format (as is output by technologies such as Illumina) and it outputs high-confidence SNP calls. SNPTB has additional scripts for data organization and management as well as downstream analyses to identify the gene(s) (or the intergenic region) that harbor the mutations and to determine if the SNP(s) in a protein-coding gene is a synonymous or a non-synonymous mutation. This pipeline is written for biologists with some exposure to the UNIX operating system.

## Implementation

SNPTB is a python package for analyzing WGS data in the fastq format. The steps in the SNPTB pipeline are: i) data organization, ii) quality control, iii) read mapping, iv) SNP calling, and v) SNP annotation (Figure 1). In the data organization step, a python script creates a new folder for each sample (prefix in the fastq filenames serves as the sample name) and moves the associated fastq files into that folder. The subsequent quality control and read mapping steps create subfolders ‘qc’ and ‘mapped’ within each sample’s directory.

**Figure 1.**
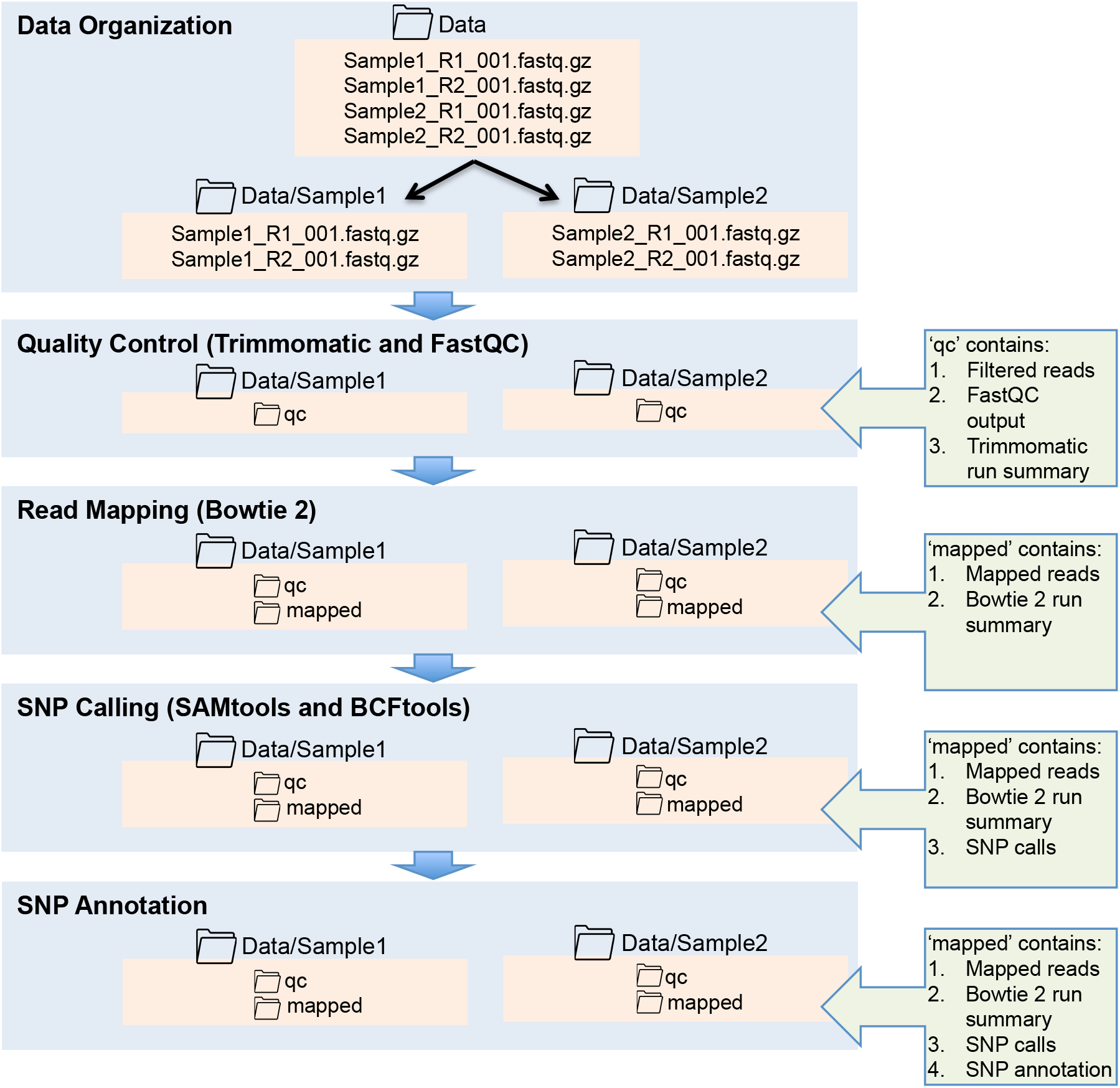
The SNPTB pipeline for WGS data analyses. Main steps in the pipeline are shown along with corresponding directory structure.

During library preparation for Illumina sequencing, short sequences called ‘adapters’ are ligated to the DNA fragments. SNPTB uses open-source software Trimmomatic (version 0.36) to remove these adapter sequences from the reads output by the Illumina sequencers and to perform quality control analysis^9^. Reads shorter than 20 nucleotides are dropped. Trimmomatic also trims low-quality ends of the reads and clips a read if the average quality score in a window of four nucleotides falls below 20. FastQC then assess the quality of the filtered high-quality reads^10^. The final filtered reads, the FastQC output, and the overview of the adapter trimming and quality control analyses are saved in the folder ‘qc’.

The high-quality filtered reads are then mapped to the *M. tuberculosis* H37Rv reference genome (NCBI Accession AL123456.3) using the open-source read aligner software Bowtie 2 (version 2.2.6)^11^. The pipeline also outputs average depth (number of reads at each genome position) for each sample as well as the percent of the genome mapped by at least 5, 10 or 20 reads. High confidence SNPs in the mapped reads (probability that a SNP call is incorrect <1e-20) are identified using open-source software SAMtools (version 1.2) and BCFtools (version 1.2)^12, 13^. The SNP calls are made relative to the *M. tuberculosis* reference genome and are further annotated to determine the location of the SNP (gene name or intergenic region) as well as the functional consequence of the mutation (synonymous or non-synonymous if the mutation lies in a protein-coding gene). The pipeline can also identify SNPs relative to another *M. tuberculosis* WGS sample instead of the reference genome.

## Discussion

While the cost of sequencing is decreasing, the high cost of computational analysis of WGS data and long turnaround time remains a challenge for biology laboratories that routinely generate appreciable amounts of sequencing data. Several open-source software for WGS data analyses exist, but they present a steep learning curve to bench biologists with limited computational experience. The goal of SNPTB is to facilitate bench biologist’s access to computational software for SNP calling, which is one of the most common and straightforward analyses of WGS data. The python package comes with an example dataset and a step-by-step tutorial to utilize SNPTB for SNP calling and SNP annotation in *M. tuberculosis* genomes.

## Acknowledgements

This work was supported by the NIH/NIAID Tuberculosis Research Unit grant U19AI11276. The funding source had no role in the study design; in the collection, analysis and interpretation of data; in the writing of the report; and in the decision to submit the article for publication.

